# Disrupting translation in CA1 pyramidal cells blocks novelty-induced increases in population firing rate while preserving newly formed assemblies

**DOI:** 10.64898/2026.11.25.769023

**Authors:** Gaokhia Yang, Nicole Maphis, Brandon T. Jackson, Matthew Bacho, Nora Perrone-Bizzozero, Sam McKenzie

## Abstract

Long-term memory depends upon the translation of the plasticity-related proteins required for synaptic consolidation. Whether learning-related changes in hippocampal CA1 coding and synchrony require local synaptic consolidation or reflect plasticity elsewhere remains unknown. We used the inducible protein kinase R (iPKR) chemogenetic system to downregulate cap-dependent translation in CaMKIIα-expressing dorsal CA1 neurons through intracerebral infusion of an actuator that activates the integrated stress response. Without chemogenetic activation, iPKR mice exhibited elevated pyramidal cell firing and reduced interneuron firing relative to wild-type mice. Infusion of the chemogenetic actuator suppressed c-Fos expression for at least 24 hours and blocked the novelty-related increase in firing observed following vehicle infusion. Nevertheless, other aspects of experience-dependent physiology persisted. Pyramidal cells active during novel exploration remained preferentially recruited into sharp-wave ripples (SPW-Rs) hours into a post-exposure consolidation period. Pairwise coincident activity patterns formed during novel exploration were also reactivated during SPW-Rs with magnitudes and temporal persistence comparable to vehicle sessions and wild-type mice without infusions. When mice returned five hours after exposure to a novel environment, the assemblies formed during translation disruption were also re-expressed at magnitudes comparable to those observed during repeated familiar-environment visits and following novel exploration under vehicle. These findings demonstrate that experience-dependent reorganization of CA1 pyramidal cell synchrony can survive disruption of plasticity-related protein production, whereas novelty-related changes in excitability are impaired. Therefore, SPW-R reactivation and cell assembly consolidation likely depend upon plasticity pathways outside of the eIF2α-dependent signaling pathways in the CA1 pyramidal cells targeted by the iPKR system.

## 1 Introduction

Experience modifies how neuronal assemblies coordinate during “online” periods of exploration when theta rhythms dominate and during “offline” moments when neurons are recruited into sharp-wave ripples (SPW-Rs) (Pavlides & Winson, 1989; Skaggs & McNaughton, 1996; Wilson & McNaughton, 1994; El-Gaby et al., 2021; Berners-Lee et al., 2022; Grosmark & Buzsáki, 2016; Silva et al., 2015; Chen et al., 2025). These changes include shifts in population firing rates (Larkin et al., 2014), preferential re-engagement of recently active neurons (Pavlides & Winson, 1989), and re-expression of ensemble co-activity for hours after a novel experience (Gupta et al., 2010; Giri et al., 2019). Although protein synthesis has been implicated in memory stabilization (Davis & Squire, 1984), it remains unclear whether the reorganization of hippocampal CA1 cell assemblies during learning requires *de novo* protein synthesis within CA1 or can be supported by synaptic plasticity in other networks.

To link synaptic plasticity to the subsequent reactivation of the CA1 assemblies that form during learning, most work has targeted the N-methyl-D-aspartate receptor (NMDAR), which is required for the induction of long-term potentiation (LTP; Harris et al., 1984; Morris et al., 1986). Systemic NMDAR antagonism during learning suppresses subsequent reactivation of newly formed assemblies during SPW-Rs(Dupret et al., 2010; Silva et al., 2015), but these drugs also reduce baseline excitatory transmission (Abraham & Mason, 1988; Hablitz & Langmoen, 1986), alter interneuron signaling (Korotkova et al., 2010), and impair learning below the concentrations that block LTP (Laha et al., 2022), confounding encoding with consolidation. CA3-restricted deletion of the NR1 subunit of the NMDAR diminishes CA1 replay of a novel experience, while leaving intact preconfigured CA1 sequences (Dragoi & Tonegawa, 2013), placing some of the required plasticity upstream, however it is unknown whether stabilization of CA1 assemblies requires local translation during encoding and consolidation.

A chemogenetic strategy has been developed to inhibit protein synthesis in a Cre-dependent and drug-inducible manner using inducible protein kinase R (iPKR) as an actuator (Shrestha et al., 2020; Figure 1A). In this system, an engineered PKR pathway promotes phosphorylation of (eIF2α), suppressing general translation initiation by 50% while engaging the integrated stress response (ISR). The antiviral drug asunaprevir (ASV) regulates this system through its interaction with the engineered NS3/4A protease that controls iPKR cleavage. Chemogenetic activation of this system has been shown to be sufficient to block LTP and long-term memory in the amygdala (Shrestha, Ayata, et al., 2020; Shrestha & Klann, 2022).

**Figure 1.**
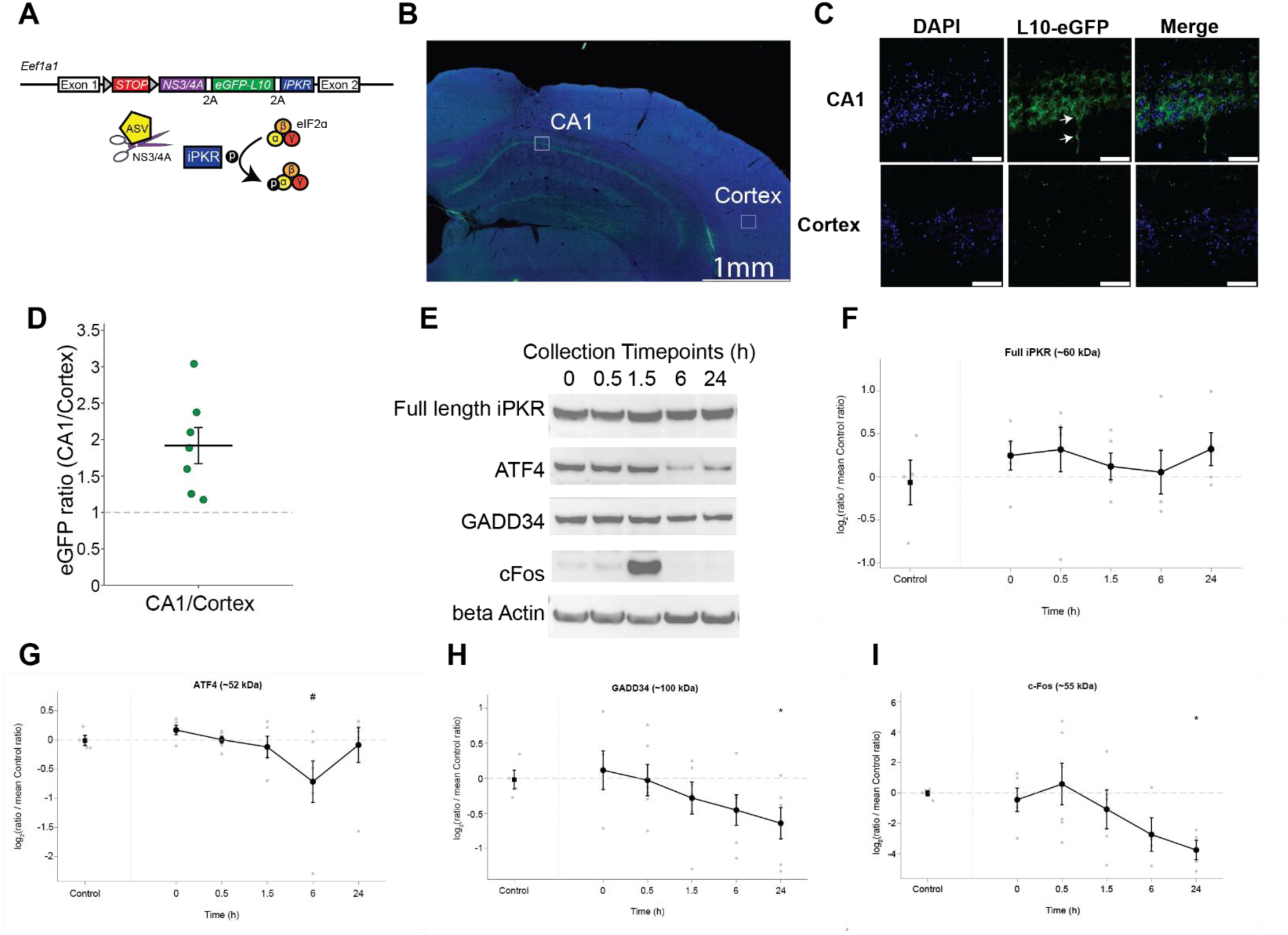
Hippocampal CA1 CaMKIIα cells activated with Cre express eGFP and undergo protein changes associated with the ISR. **A**, The iPKR system is a transgenic mouse model with a multicistronic cassette between exon 1 and exon 2 of the eukaryotic elongation factor 1a1 (Eef1a1) gene. The cassette contains transcripts for an engineered protein kinase R (iPKR) that harbor a recognition site for a recombinase non-structural protein 3/4 (NS3/4A) protease. ASV administration inhibits NS3/4A protease activity, permitting iPKR to phosphorylate eIF2α and initiating global protein translation or cap-dependent protein inhibition and activating the ISR or cap-independent protein translation. The L10-eGFP allows for ribosomal tagging of the large subunit with a green fluorescent protein. The self-cleaving 2A proteinase sites allow for ribosomal skipping during transcription (Shrestha, Ayata, et al., 2020). **B,** An injected mouse showed the expected eGFP expression in the pyramidal layer highlighted in the CA1 white box at 20 X objective. **C,** Insets at 63X showing the non-transfected cortex of an hippocampally-infused subject to normalize background fluorescence. The white arrows indicate the eGFP expression in the soma (top arrow) and projection (bottom arrow). The white scale bar represents 25 µm. **D,** Mean ratio comparing the fluorescence of eGFP to DAPI between the CA1 pyramidal layer to cortex (n = 7 electrophysiology mice, mean ± SEM) **E,** Representative Western blot of individual subjects sacrificed per each time point. **F,** The full-length iPKR construct was stable across 24 hours in experimental animals and was similar to controls (Welch’s t-test Control vs 0: p=0.342, Control vs 0.5: p=0.204, Control vs 1.5: p=0.603, Control vs 6: p=0.703, Control vs 24: p=0.274; 0 vs 0.5: p=0.615, 0 vs 1.5: p=0.573, 0 vs 6: p=0.686, 0 vs 24: p=0.737). **G,** ATF4 significantly decreased at 6 h post-ASV infusion compared to timepoint 0 but not controls (Welch’s t-test Control vs 0: p=0.171, Control vs 0.5: p=0.916, Control vs 1.5: p=0.716, Control vs 6: p=0.073, Control vs 24: p=0.884; 0 vs 0.5: p=0.131, 0 vs 1.5: p=0.208, 0 vs 6: p=0.019, 0 vs 24: p=0.498). **H,** GADD34 steadily declined over time and significantly was lower than control at 24 hours (Welch’s t-test Control vs 0: p=0.528, Control vs 0.5: p=0.832, Control vs 1.5: p=0.407, Control vs 6: p=0.178, Control vs 24: p=0.050; 0 vs 0.5: p=0.676, 0 vs 1.5: p=0.283, 0 vs 6: p=0.171, 0 vs 24: p=0.097). **I,** c-Fos steadily declines 0.5 h after infusion and is significantly lower than control at 24 h (Welch’s t-test Control vs 0: p=0.773, Control vs 0.5: p=0.196, Control vs 1.5: p=0.605, Control vs 6: p=0.129, Control vs 24: p=0.002; 0 vs 0.5: p=0.205, 0 vs 1.5: p=0.690, 0 vs 6: p=0.202, 0 vs 24: p=0.078). * Indicates p-value < 0.05 compared to control. # Indicates p-value < 0.05 compared to timepoint 0.

Here, we used the iPKR system to test whether cell-intrinsic protein synthesis is required to stabilize the CA1 cell assemblies formed during novel context exploration. We found that activating the iPKR system reduced production of the immediate early gene c-Fos for at least 24 hours, and blocked the normal increase in population firing rate after exposure to a novel environment. Reactivation of assemblies into SPW-Rs and during subsequent context re-exposure was comparable to that observed during within-subject control sessions involving saline infusion. The findings support a model in which CA1 pyramidal cell plasticity drives changes in excitability following novelty, but not the fine-time scale reorganization of CA1 assemblies, which likely depends upon plasticity in other circuits.

## 2 Results

### Activation of the iPKR alters translation of proteins in the ISR and the plasticity-related protein c-Fos

To control translation in CA1 pyramidal cells, the iPKR construct was driven by AAV5-CaMKIIα-Cre infused unilaterally or bilaterally into dorsal CA1 (Figure 1A). Neurons transduced with the virus expressed L10-eGFP reporter in somata and projections (Figure 1B-D; eGFP to DAPI mean ratio = 1.92±0.25), confirming expression in CA1. The iPKR system should enable ASV to activate the ISR in the transduced CaMKIIα+ neurons; therefore, we tested for the abundance of ISR proteins ATF4 and GADD34 in hippocampal lysates collected at 0, 0.5, 1.5, 6, and 24-hr after ASV, and in control subjects (Figure 1E). We measured the iPKR full length fragment that includes the full construct and confirmed that the expression remained stable after ASV infusion, as expected (Figure 1F). In contrast to the original description of the iPKR system, ATF4 showed a significant decrease in protein levels at 6-hrs post-infusion compared to timepoint 0 but not control (Figure 1G; 0 vs 6-hr: p=0.019; Control vs 6-hr: p=0.073) and not at other collection time points (all p > 0.10). Levels of GADD34, the inhibitory feedback arm of the ISR (Novoa et al., 2001; Oliveira et al., 2024), also steadily decreased, reaching its lowest point 24-hours after ASV infusion (Figure 1H); GADD34 levels in iPKR mice showed a non-significant elevation relative to control mice (Control vs 0-hr: p=0.528) and steadily declined out to 24 hours with significance compared to control (Control vs 24-hr: p=0.050). Finally, we measured c-Fos, a short-lived, plasticity-related immediate early gene product whose expression is suppressed by activation of the integrated stress response (Shrestha et al., 2020). c-Fos protein levels were initially elevated, and then reduced at 6-hrs after infusion (Figure 1I; Control vs 6-hr: p=0.129). The suppression was still measured 24 hours post ASV infusion (0-hr vs 24-hr, p=0.078; Control vs 24-hr, p=0.002), extending the duration of translational disruption from that reported in the original description of the iPKR system that looked out to 6 hours (Shrestha, Ayata, et al., 2020).

### Baseline firing differs between iPKR and never-infused mice

To measure changes in assembly reactivation, single units were recorded with silicon probes directed towards dorsal CA1 (Supplemental Figure 2) alongside infusion cannula implanted into the ipsilateral lateral ventricle. Outside of infusion periods, the iPKR cohort exhibited elevated CA1 pyramidal firing relative to never-infused reference mice (Huszár et al., 2022; mean firing rate = 1.60 Hz versus 0.77 Hz; mixed-effects *p* = 0.026), whereas interneuron firing was reduced (mean firing rate = 4.14 Hz versus 7.62 Hz; permutation *p* = 0.029, mixed-effects *p* = 0.072; Figure S3A). The pyramidal elevation was present in NREM (iPKR mean firing rate = 1.69 Hz versus control rate = 0.72 Hz; *p* = 0.002) and wake (1.48 versus 0.75 Hz; *p* = 0.046), while interneuron reduction was mostly confined to wake (iPKR mean firing rate = 4.06 Hz versus control rate = 9.04 Hz; *p* = 0.005; NREM, *p* = 0.10; Figure S3B). Pyramidal firing within ripples was comparable between cohorts (iPKR mean firing rate = 4.11 versus control rate = 4.27 Hz; *p* = 0.71), resulting in substantially lower ripple-associated gain in iPKR mice (iPKR = 2.41 versus control = 6.44; *p* = 0.005; Figure S3C). Ripple duration and peak frequency did not differ (*p* = 0.33 and *p* = 0.91; Figure S3D).

### ASV blocked novelty-associated firing rate increases in CA1 during NREM

To test how chemogenetic disruption of protein translation affects neurophysiological correlates of memory, we exposed mice to familiar (Familiar 1) and novel (Novel 1) testing environments and monitored the stability of neural coordination during SPW-Rs recorded in the home cage during a 4-hr consolidation period, and during context re-exposure (Familiar 2 and Novel 2; Figure 2A). Two controls were built into the experimental design. First, mice were exposed to a familiar environment prior to drug infusion; based on previous studies (Agnihotri et al., 2004), we predicted that place fields would remain stable and that replay of these previously consolidated maps would persist despite disruption of synaptic consolidation. Second, we employed a within-subject control in which saline (SAL) vehicle was infused instead of ASV. Due to concerns about long-term effects of ASV, most SAL sessions were conducted prior to ASV (SAL before ASV in N = 9 animals; ASV before SAL in N = 1; N = 2 received ASV only). Of 16 SAL sessions, 13 preceded the animal’s first ASV session.

**Figure 2.**
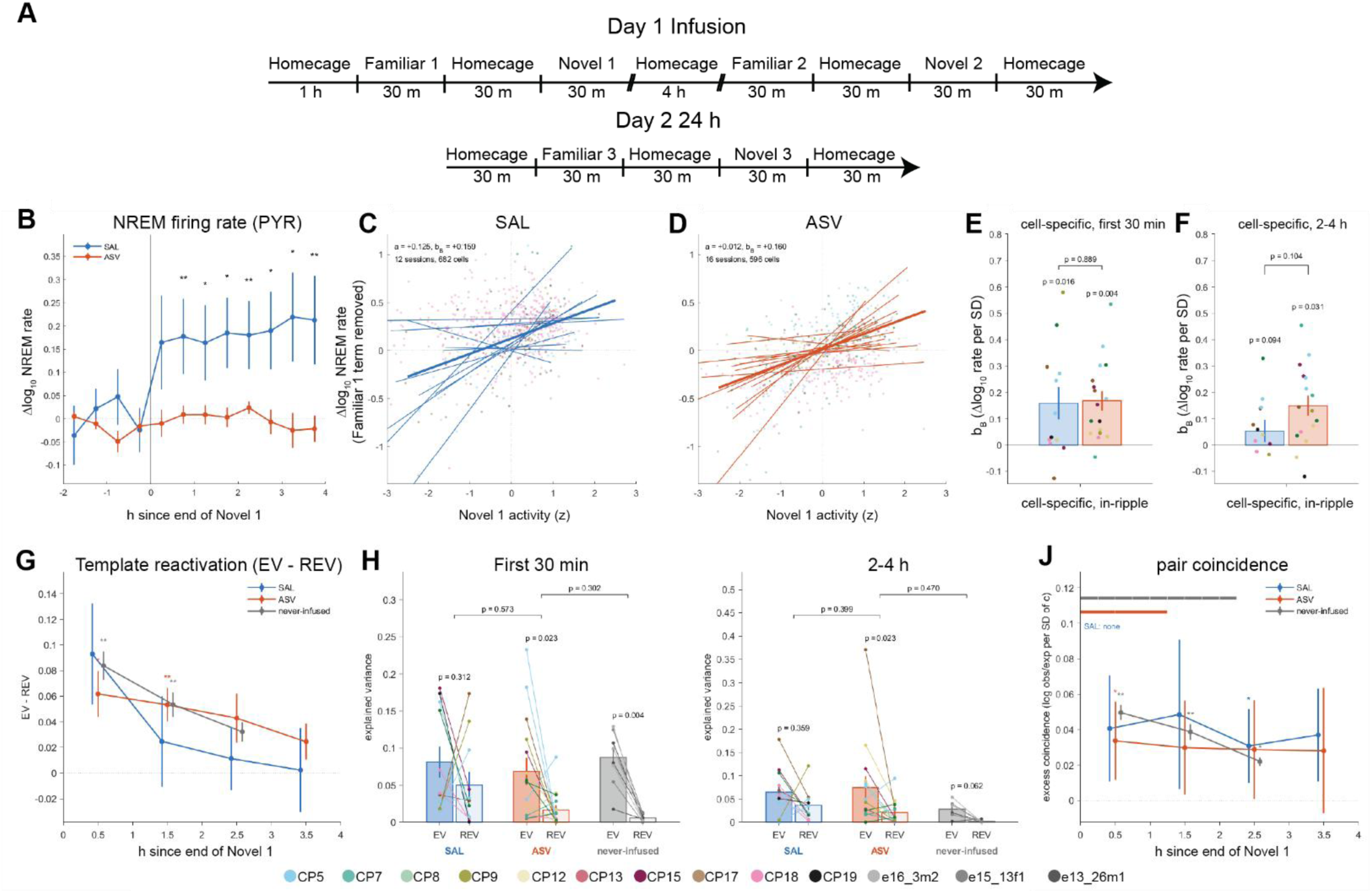
ASV activation blocks changes in CA1 firing rate after novelty, but preserved the reactivation of assemblies into SPW-R. **A**, Recording schedule. On day 1, mice were recorded in the home cage and exposed for 30 min each to a familiar (Familiar 1) and a novel (Novel 1) context, then returned to both contexts after 4 h (Familiar 2, Novel 2); all contexts were revisited on day 2 (Familiar 3, Novel 3). Saline (SAL) or ASV was infused in the home cage between Familiar 1 and Novel 1. **B**, Change in NREM firing rate of pyramidal cells, expressed as the change in log10 rate relative to before the infusion, in 30-min bins aligned to the end of Novel 1 (mean ± SEM across animals; SAL, 14 sessions from 9 animals; ASV, 16 sessions from 10 animals). Asterisks mark bins in which SAL and ASV differ (paired sign-flip test across animals). **C, D**, Partial-residual plots of the change in log10 in-ripple firing rate from the 30 min before Novel 1 to the first 30 min after it, with the familiar-context term removed, against each cell’s activity during Novel 1 (z-scored within session), for SAL (C; 682 cells, 12 sessions) and ASV (D; 596 cells, 16 sessions). Dots are cells colored by mouse, thin lines are per-session fits, and the thick line is the mean fit. The intercept of each fit is the change shared by all cells and the slope is the cell-specific component. **E, F**, Cell-specific component (slope) per session in the first 30 min (E; SAL, 12 sessions from 8 animals; ASV, 17 sessions from 9 animals) and 2–4 h (F; SAL, 12 sessions from 8 animals; ASV, 16 sessions from 9 animals) after Novel 1. One ASV session lying more than 7 robust SD from the group median was excluded from F. **G**, Template reactivation (explained variance minus reverse explained variance, EV − REV) in 1-h bins after Novel 1, with ripple numbers equated across bins by subsampling (mean ± SEM across animals; first bin: SAL, 11 sessions from 7 animals; ASV, 15 sessions from 8 animals; never-infused, 9 sessions from 3 animals). Asterisks mark EV − REV greater than zero (sign-flip test across animals, or across sessions for never-infused mice; uncorrected). **H, I**, EV and REV per session in the first 30 min (H) and 2–4 h (I) after Novel 1, with lines joining each session’s values (SAL, 11 sessions; ASV, 15 sessions; never-infused, 9 sessions in H and 6 in I). p values above each group test whether EV − REV is greater than zero; brackets compare EV − REV between groups (linear mixed-effects model with a random intercept for animal). **J**, Excess coincident firing during ripples of cell pairs correlated during Novel 1, in 1-h bins after Novel 1, expressed as log(observed/expected coincidences) per SD of the exploration pair correlation (mean ± SEM across animals; SAL, 12 sessions from 8 animals; ASV, 18 sessions from 10 animals; never-infused, 9 sessions from 3 animals). Asterisks as in G. Bars along the top mark windows in which the excess remained significant in a 15-min scan, with p values calibrated for window selection (p < 0.05). Bars in E, F, H, and I show mean ± SEM across sessions; p values above bars test for a difference from zero (sign-flip test across animal means); brackets in E and F compare groups by permutation across sessions. Dots are colored by mouse (key at bottom).

Following novel context exposure (>2 hours after infusion), pyramidal cell NREM firing rose after a first saline infusion from 2.00 ± 0.40 to 2.84 ± 0.45 Hz (+0.185 ± 0.110 log10 units, ×1.53; N = 8 mice) and was unchanged across ASV sessions (2.01 ± 0.22 to 1.96 ± 0.24 Hz; −0.007 ± 0.030, ×0.98; N = 12 mice; ASV – saline: −0.207 ± 0.094, p = 0.036; Figure 2B). Interneuron firing changed in the same direction (saline +0.134 ± 0.076; ASV −0.009 ± 0.050), but the arms did not differ significantly (order-adjusted −0.227 ± 0.139, p = 0.11). Between the infusion and the novel context exposure, firing rate did not change significantly, and the ASV vs SAL sessions were indistinguishable (pyramidal, ASV minus saline −0.017, *p* = 0.16; *p* = 0.34; interneurons −0.000, *p* = 1.00), suggesting that the cause of the rate increase was contextual novelty, not the infusion protocol. Infusion order did not account for the contrast. Repeat saline infusions changed pyramidal firing as much as first infusions (+0.223 ± 0.204 versus +0.150 ± 0.122; difference +0.072 ± 0.237, *p* = 0.73). Repeated ASV sessions, meaning an ASV session preceded by ASV in the same animal, showed a similar decline in whole-recording pyramidal firing (−0.057 ± 0.019 log_10_units, × 0.88; ten sessions, seven animals; sign-flip against 0, *p* = 0.016). Thus, ASV blocked a novelty-associated rate increase in CA1 firing during NREM.

### ASV does not disrupt recruitment of neurons that were active in a novel experience into SPW-Rs

Classic studies of replay find that the cells that are most active during an experience tend to reactive during subsequent sleep (Pavlides & Winson, 1989). Since we observed a general rate increase after novelty, we first tested whether ripple firing was also elevated. Without considering which cells were active during novelty, we found that pyramidal ripple participation rose after saline (0.243 ± 0.027 to 0.266 ± 0.029) and fell after ASV (0.189 ± 0.016 to 0.165 ± 0.018; paired difference in differences p = 0.016, seven animals). In-ripple firing also rose after saline (5.23 to 6.95 Hz; log₁₀ 0.719 ± 0.073 to 0.842 ± 0.124) and fell after ASV (4.05 to 3.75 Hz; 0.608 ± 0.065 to 0.574 ± 0.090; p = 0.062). The ratio of in-ripple to out-of-ripple firing within the same window fell after saline (2.76 to 2.23) and rose after ASV (2.23 to 2.44), but the change did not differ between arms (p = 0.45). The increase in ripple recruitment therefore followed background firing rather than a selective change in ripple-related gain.

To distinguish this population-wide rate modulation from experience-associated changes, the change in each neuron’s activity between the home-cage periods before and after the novel exposure was decomposed into a shared term (common to all neurons and reflecting a global change in activity), and a cell-specific term (predicted by the neuron’s firing during novel-context exploration). Two aspects of ripple activity were examined: in-ripple firing rate, the number of spikes a neuron fired per second of ripple time, and ripple participation, the fraction of ripples in which it fired at least once.

Within 30 minutes of the novel context, the shared change in log in-ripple firing was positive after SAL (0.125 ± 0.044), consistent with the rise in firing after SAL infusion, and near zero after ASV (–0.005 ± 0.024; permutation p = 0.008). The same held for out-of-ripple firing (0.126 ± 0.041 versus –0.002 ± 0.025; permutation p = 0.003), suggesting that the shared response under ASV reflects a broader change in excitability rather than a cell-specific change in ripple recruitment.

In contrast, the term relating in-ripple firing to prior novel-context activity was similar for saline and ASV. The neurons that fired more during novel-context exploration increased their in-ripple firing after SAL (0.159 ± 0.061 log units per standard deviation of novel-context activity, 12 sessions; Figure 2C) and ASV (0.168 ± 0.036, 17 sessions; between-arm p = 0.90; Figure 2D). The cell-specific term capturing reactivation of recently active neurons was above zero in both arms (saline, p = 0.016; ASV, p = 0.004; sign-flip over animals; Figure 2E). In addition to changes in ripple rate, novel-context activity likewise predicted an increase in ripple participation (controlling for out-of-ripple rate) by a similar amount in both arms (fraction of ripples per standard deviation of novel-context activity: SAL, 0.020 ± 0.011; ASV, 0.024 ± 0.008; between-arm permutation p = 0.74, paired p = 0.85, eight animals) and in never-infused recordings (0.020 ± 0.003). This coefficient was above zero after ASV (p = 0.012) but less consistent across SAL animals (p = 0.22).

To control for non-specific drift, the same model was fitted to a placebo interval, from before the first familiar context exposure to before the first novel context exposure. The real-minus-placebo contrast for the cell-specific in-ripple term was positive after saline (+0.181, p = 0.034, 11 sessions) and after ASV (+0.150, p = 0.001, 17 sessions), indicating that the changes in in-ripple firing reflect activity in the novel context rather than drift. In never-infused recordings the contrast was comparable to that observed in the ASV and SAL sessions (0.13, positive in all nine sessions, p = 0.004).

To test whether these changes persisted beyond 30 minutes, the decomposition was repeated 2–4 h after the novel exposure (Figure 2F). The shared in-ripple term remained elevated after SAL and absent after ASV (0.142 ± 0.046 versus 0.025 ± 0.019; permutation p = 0.013). The cell-specific in-ripple term remained detectable after ASV (0.150 ± 0.038, 16 sessions, p = 0.031) but not after saline (0.053 ± 0.042, p = 0.094); the arms did not differ (between-arm p = 0.10). These observations indicate that experience-associated cell-to-cell differences in subsequent activity remain detectable under ASV for at least 2–4 h, while the global changes in firing rate are blocked.

### Patterns of neuronal co-activity formed during a novel experience are reinstated into SPW-Rs after ASV

Next, we tested whether ASV prevents the patterns of activity observed in a novel context from being re-expressed in subsequent ripples. First, we used the standard explained variance (EV) and reverse explained variance (REV) measures to assess reactivation during post-exploration ripples. EV quantifies how much of the variance in pairwise spike-count correlations during post-exploration ripples is explained by correlations during exploration, after accounting for correlations during pre-exploration ripples. REV exchanges the pre– and post-exploration ripple periods, providing a baseline for the contribution of pre-existing correlation structure. In SAL sessions, EV and REV were both elevated (0.081 ± 0.021 and 0.050 ± 0.018, respectively; EV − REV, 0.031 ± 0.034, p = 0.31), indicating that pre-exposure ripple coactivity already resembled the subsequent novel-context template. This limits the utility of the raw saline EV − REV value as a treatment comparator (Figure 2G). ASV sessions were therefore compared with never-infused recordings (Huszár et al., 2022). In the first 30 min after exploration, EV − REV after ASV was 0.052 ± 0.021 (15 sessions, eight animals), positive in seven of eight animals (p = 0.023), and did not differ detectably from never-infused recordings (0.082 ± 0.011; mixed-effects p = 0.30; Figure 2H). EV – REV after ASV declined gradually over the following hours (Figure 2G) and remained positive when pooled over 2–4 h after the exposure (0.054 ± 0.024, p = 0.023; Fig. 2I), reflecting a similar time course as that observed in the single-neural reactivation analysis (Figure 2F).

EV−REV does not by itself show that pairwise patterns are reinstated, since a cell-autonomous rule that allowed recently active neurons to be recruited into ripples would likewise drive increase EV−REV without any pair-specific structure. We therefore modeled spike coincidences between pairs of pyramidal neurons in individual ripples while regressing out baseline pairwise structure and a slowly-varying marginal ripple firing rate (Materials and Methods). In never-infused recordings, pairs that were more correlated during the novel exposure co-fired more than expected in the ripples that followed it. In the first hour, coincidences exceeded expectation by 5.1 ± 0.5% per standard deviation of the exploration pair correlation (9 sessions from three mice; p = 0.004) which decayed to 2.2 ± 0.2% in the third (six sessions with recordings that long; p = 0.031; Figure 2J). These levels of excess synchrony matched those observed following ASV or SAL (mixed-effects p ≥ 0.66). After ASV it was 3.4 ± 2.3% in the first hour after the novel exposure (18 sessions, ten animals; p = 0.014) versus 4.2 ± 3.1% in SAL (sign-flip test against 0, p= 0.07); the arms did not differ (mixed-effects p = 0.36). To ask how long the excess remained detectable at finer resolution, ripples were grouped into 15-min bins after the exposure. Neighboring bins were pooled until the excess became significant, and the p-value was recalibrated for this pooling by repeating the procedure on data with the sign of each session’s effect randomly flipped. The novel-exposure excess after ASV remained detectable in the window 0.50–1.25 h after the novel context exposure (calibrated p = 0.042). After SAL, no window reached calibrated significance (0–1.25 h, calibrated p = 0.098), although the hourly excess after saline was at least as large as after ASV. In never exposed subjects, the novel-exposure excess remained detectable in the window 2.00–2.25 h after the exposure (calibrated p = 0.016; six sessions with recordings that long). A control home-cage template showed no excess in either arm in any hour (p ≥ 0.12).

Thus, pair-specific reinstatement of exploration co-activity was detectable for at least 2-hrs in never-infused recordings, and an excess of comparable size was present in every hour after either infusion. The infused recordings did not resolve when the excess ended in either arm, so they cannot show whether persistence differs under ASV, though predictable covariance structure was observed >30 minutes after the novel context exposure despite translation disruption by ASV, and single cell changes in ripple recruitment were detectable for at least 2-hrs after novel context exposure.

### Novelty-related patterns of activity formed under ASV are re-expressed hours later upon return to the encoding environment

In each session, mice revisited the familiar environment (Familiar 2) 237.9 ± 16.8 min after the first novel exposure (Novel 1) and the novel environment (Novel 2) 329.7 ± 10.6 min after Novel 1. The Novel 1– Novel 2 interval spans a period during which long-term memory and long-term potentiation are thought to depend on the synthesis of plasticity-related proteins (Fonseca et al., 2006; Remaud et al., 2014; Rosenberg et al., 2014). In ASV sessions, Novel 1, Familiar 2 and Novel 2 all followed the infusion, so coordination in the novel environment was first expressed, and later re-expressed, with ASV on board. For each exposure, the coordination of every pair of pyramidal neurons was measured as the partial correlation of their spike counts in 100-ms bins during running, with the summed count of the other neurons partialled out to account for changes in population firing rate (Materials and Methods).

Pairwise coordination in Novel 2 resembled that in Novel 1 more than that in the Familiar 1 (Figure 3A). The patterns of pairwise coincidence in Novel 1 explained those observed in Novel 2 to a greater degree than patterns in Familiar 1 for ASV (Novel 1>Familiar 1 as a predictor of Novel 2 = 0.134 ± 0.042, 17 sessions, 10 mice; p = 0.016) and SAL sessions (Novel 1>Familiar 1 as a predictor of Novel 2 = 0.177 ± 0.101, 13 sessions, 8 mice; p = 0.13; Figure 3B). Because Novel 1 was closer in time to Novel 2 than Familiar 1 was, this comparison favors Novel 1 for reasons of elapsed time alone, however, controlling for elapsed time did not change the explanatory power of Novel 1 assemblies in explaining Novel 2 patterns of coincidence (Novel 1>Familiar 1 as a predictor of Novel 2 = 0.134 ± 0.041 after ASV, p = 0.008; Novel 1>Familiar 1 as a predictor of Novel 2 = 0.159 ± 0.090 after SAL, p = 0.13) and there was no effect of ASV treatment on the specificity of pairwise reactivation within the novel context (paired difference –0.015 ± 0.071, p = 0.87, eight animals; mixed-effects with recording day as a covariate, p = 0.92; unadjusted comparison –0.022, p = 0.73; Figure 3C).

**Figure 3.**
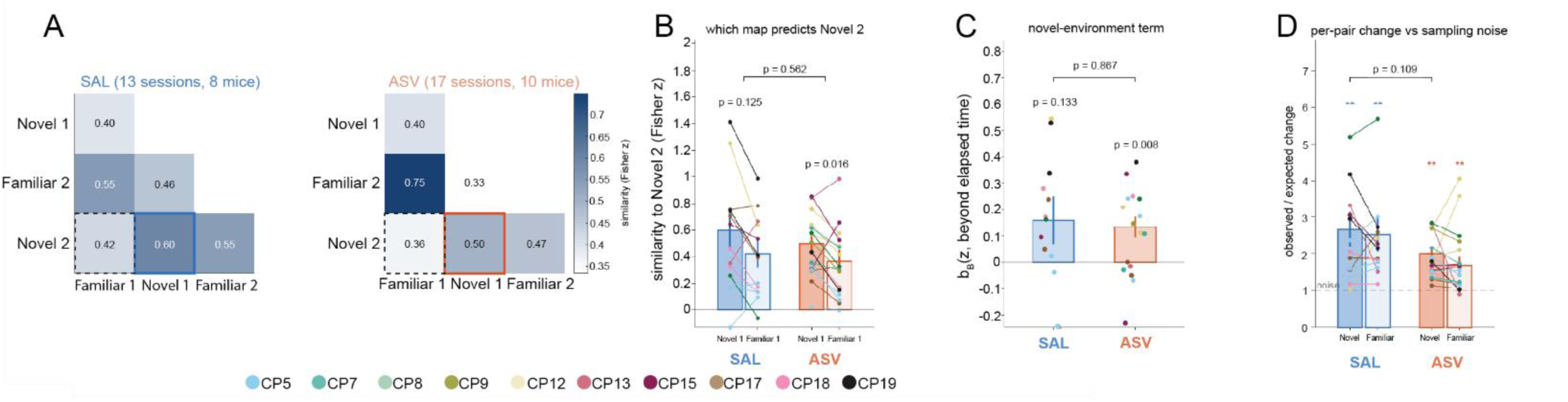
Coordination patterns formed in the novel context are re-expressed on return under saline and ASV. **A**, Similarity of coordination patterns between exposures for SAL (13 sessions from 8 mice) and ASV (17 sessions from 10 mice), expressed as the Fisher z-transformed correlation, across cell pairs, of the pairwise spike-count correlations measured in each exposure (Familiar 1; Novel 1; Familiar 2; Novel 2). Values are means across animals; solid outlines mark the Novel 1 to Novel 2 comparison and dashed outlines the Familiar 1 to Novel 2 comparison. **B**, Similarity of Novel 2 to Novel 1 and to Familiar 1 per session, with lines joining each session’s two values. p values above each group test whether similarity to Novel 1 exceeds similarity to Familiar 1; the bracket compares this difference between SAL and ASV (paired test across the 8 animals recorded under both). **C**, Novel-environment term per session: the similarity between Novel 1 and Novel 2 beyond that expected from the time elapsed between them, estimated from a regression of all six pairwise similarities on the time between exposures, with separate terms for familiar–familiar and novel–novel pairs. p values above bars test for a difference from zero; the bracket compares SAL and ASV as in B. **D**, Change in individual pair correlations between exposures relative to sampling noise, for Novel 1 to Novel 2 and Familiar 1 to Familiar 2, computed as the median absolute change in pair correlation divided by the change expected from sampling noise alone. The dashed line marks a ratio of 1 (change within noise), asterisks mark ratios greater than 1, and the bracket compares the Novel 1 to Novel 2 ratio between SAL and ASV as in B. In B–D, bars show mean ± SEM across animals and dots are sessions colored by mouse (key at bottom); p values and asterisks are from sign-flip tests across animal means (**p < 0.01).

Pairwise coordination of assemblies formed under ASV was no less stable than those measured in the familiar environment, nor less stable than under SAL. Stability was measured pair by pair, as the median change in a pair’s coordination between two exposures relative to the change expected from sampling noise alone (Figure 3D). Pairwise coordination decreased in both contexts across the 4-5 hours period, as expected from representational drift (Khatib et al., 2023). However, under ASV the novel map did not change detectably more between Novel 1 and Novel 2 than the familiar map did between Familiar 1 and Familiar 2 (2.0 ± 0.2 versus 1.7 ± 0.2 times the noise level, difference 0.32 ± 0.24, p = 0.24, N = 10 mice); nor was the novel map less stable after ASV than after SAL. If anything, its change was smaller after ASV (by 0.72 ± 0.39; p = 0.11, N = 8 paired mice; mixed-effects p = 0.034), as was the familiar map’s (by 0.90 ± 0.40; p = 0.047; mixed-effects p = 0.011). Thus, pairwise coordination expressed in a novel environment under ASV was preferentially re-expressed on return to that environment about 5.5-hrs later, to a degree not detectably different from that after saline.

## Discussion

Acute activation of the iPKR system in CA1 CaMKIIα+ neurons had separable effects on neuronal responses to a novel environment. The rise in NREM population firing rate that followed the novel exposure after saline, was absent following chemogenetic disruption of CA1 protein translation. On saline days, firing rate was unchanged between the infusion and the novel exposure, so the rise followed the experience rather than the infusion. In contrast, experience-related reactivation of patterns of neuronal co-activity during SPW-Rs remained detectable after ASV at levels comparable to those after control infusions. Likewise, patterns of neuronal coordination formed in a novel environment were re-expressed after ASV when mice returned hours later. These results support a distinction between experience-dependent changes in population firing rate, which depended on normal protein translation in CA1, and the experience-related changes in pairwise coactivity patterns expressed during SPW-Rs and on return to the novel environment, which were relatively spared.

### Novelty-associated increases in CA1 firing blocked by activating the ISR

Novel context exposure normally elevates CA1 population firing, including during subsequent ripples (Karlsson & Frank, 2008; Larkin et al., 2014; Hirase et al., 2001). ASV abolished that increase (Figure 2B), suggesting a mechanism that is intrinsic to CA1 pyramidal cells. One possibility is that the rate increase is a downstream consequence of cap-dependent translation inhibition, in which case its loss reflects blocked consolidation at the synapse (Chen et al., 2017) or even at the level of cell-intrinsic excitability (Cohen-Matsliah et al., 2010). A second possibility, is that eIF2α-dependent signaling suppresses excitability more directly, through routes that do not require bulk translational arrest. For example, through interferon-γ-mediated disinhibition (Zhu et al., 2011), and local PERK reduction increases CA1 excitability (Sharma et al., 2018). This second possibility is given weight by work showing that enhancing eIF2α phosphorylation in CA1 pyramidal cells impairs consolidation while elevating ATF4 and suppressing CREB-dependent signaling, without an evident reduction in general translation (Jiang et al., 2010), and the timing of the ASV effects on firing rate in our own experiments, which emerged prior to decreases in protein expression.

Activation of iPKR produced a biphasic change in c-Fos protein: a transient early rise followed by suppression that persisted 24 hours after ASV infusion (Figure 1I). Because all control tissue was collected prior to the expected windows in which c-Fos is expressed after sustained neuronal activity (Zhang et al., 2002), we cannot distinguish whether the early rise reflects the infusion itself or the acute stress-kinase signaling that drives immediate early gene induction after translational perturbation (Edwards & Mahadevan, 1992; Iordanov et al., 1997; Cano et al., 1994). The direction of the change between ATF4 and GADD34 also declined, with the GADD34 nadir lagging that of ATF4, as expected given that GADD34 induction is transcriptionally dependent on ATF4 (Ma & Hendershot, 2003). The direction of the ATF4 change is nonetheless opposite to the prediction of canonical integrated stress response signaling, in which eIF2α phosphorylation represses bulk translation while selectively increasing the translation of ATF4 (Harding et al., 2000; Ma & Hendershot, 2003; Jiang et al., 2010). A coordinated decrease in several short-lived proteins, ATF4 among them, is better described as broad suppression of *de novo* translation in which the privileged translation of ATF4 is not sustained, consistent with the original characterization of this system (Shrestha, Ayata, et al., 2020). Therefore, progressive loss of GADD34, which normally terminates the response, offers a candidate explanation for the persistence of suppression (Hayakawa-Ogura et al., 2023).

We cannot account for the baseline firing rate differences in iPKR mice in the absence of ASV. Sustained manipulation of excitability-related transcriptional pathways can itself damage hippocampal neurons (Lopez De Armentia et al., 2007) and a convulsive seizure was observed in a non-implanted subject.

### Synaptic consolidation, place field stability, and the separability of rate from ensemble structure

We observed intact reactivation of CA1 assemblies formed in a novel environment under ASV, both during post-exposure SPW-Rs and upon return to that environment five hours later. Manipulations of plasticity signaling have been shown to disrupt CA1 coding and assembly coordination. Genetic disruption of transcription-dependent plasticity destabilizes the long-term CA1 spatial code (PKA: Rotenberg et al., 2000; CREB: Cho et al., 1998; zif268/egr1: Renaudineau et al., 2009; Fos: Pettit et al., 2022; DNA methylation: Roth et al., 2015), as does interference with induction and receptor-delivery machinery (αCaMKII: Cho et al., 1998; Rotenberg et al., 1996; CA1 NMDAR: McHugh et al., 1996; CA1 Syntaxin3: Plitt et al., 2026). Reversing structural LTP by light-inactivation of cofilin in CA1 CaMKIIα neurons, either immediately after encoding or throughout the ensuing sleep, prevents the next-day synchronization of cells coding a shock experience (Goto et al., 2021). Systemic protein synthesis inhibition with anisomycin leaves place field stability intact at one hour while disrupting it six hours later (Agnihotri et al., 2004), about the interval separating our two exposures. Systemic NMDAR antagonism likewise disrupts reactivation of recently formed assemblies within the first hour after learning, when our ripple measurements were made (Dupret et al., 2010; Silva et al., 2015). Our observation of preserved reactivation is therefore unlikely to reflect measurement at too short a delay for plasticity-dependent destabilization to emerge, nor an ineffective manipulation, since ASV blocked the novelty-induced rate increase in the same animals. Fine-timescale synchrony and its re-expression appear not to require the signaling engaged by iPKR activation in CA1 CaMKIIα neurons.

Activation of the ISR left baseline activity intact, unlike translation elongation inhibitors, which profoundly suppress local field potentials and multiunit firing to a degree comparable with inactivation (Sharma et al., 2012; Scavuzzo et al., 2019; Al-Smadi et al., 2024). At the single-cell level, brief anisomycin exposure lowers firing frequency, reduces spike amplitude and broadens spikes across a train while leaving input resistance, membrane time constant and spike threshold unchanged — a profile pointing to failure of ionic gradient maintenance rather than to loss of any plasticity-related protein (Scavuzzo et al., 2019). Emetine behaves similarly (Al-Smadi et al., 2024), and differential effects of puromycin and cycloheximide on hippocampal electrical activity were reported more than half a century ago (Cohen et al., 1966). Because these agents perturb excitability and metabolism alongside translation, they cannot distinguish lost plasticity from lost firing; the present approach separates the two, and shows that the novelty-driven rate component and the ripple covariance structure have different requirements for eIF2α-dependent signaling in CA1 pyramidal cells.

We observed that pairwise firing relationships were more stable with ASV infusions, suggestive of a disruption in the plasticity processes driving decorrelation related to representational drift. Studies on representational drift dissociate single-cell field persistence and population-level representational stability and it is increasingly understood that representational drift may require experience-dependent plasticity (Ratzon et al., 2024). CA1 shows substantial cell-level turnover across days while the spatial and contextual code remains decodable (Ziv et al., 2013; Keinath et al., 2022; Gonzalez et al., 2019), contextual representation is preserved despite ongoing drift (Keinath et al., 2022). Firing rate and tuning are likewise separable axes of drift rather than a single quantity (Abbaspoor et al., 2026; Geva et al., 2023; Khatib et al., 2023; Tang et al., 2026). Related dissociations appear within ripples themselves: experience-driven rate modulation is reinstated during replay (Tirole et al., 2022), and slow-firing, experience-responsive cells gain ripple recruitment independently of population rate shifts (Grosmark & Buzsáki, 2016).

If CA1-local translation is not required for the reactivation we measured, the plasticity supporting it may reside elsewhere. CA3 output is required for ripple-associated reactivation and for consolidation (Nakashiba et al., 2009), and CA3 synaptic plasticity specifically supports ripple physiology during consolidation (El Oussini et al., 2023); CA3-restricted NMDAR deletion strengthens preplay relative to replay, consistent with a block on the plasticity that would otherwise incorporate novel information into subsequent ripples (Dragoi & Tonegawa, 2013). CA1 assembly content is further shaped by medial entorhinal input (Zutshi & Buzsáki, 2023; Yamamoto & Tonegawa, 2017), by CA2 (He et al., 2021), and by dentate circuitry (GoodSmith et al., 2025). Alternatively, replay may depend on plasticity in inhibitory cells. Inhibitory neurons are themselves loci of translation for long-term memory under this same chemogenetic strategy (Shrestha, Shan, et al., 2020) and eIF2α-dependent consolidation has been localized to somatostatin neurons (Sharma et al., 2020). Learning reconfigures pyramidal–interneuron coupling in CA1, which has been related to changes in replay content (Dupret et al., 2013). Since interneuron populations gate network stabilization and ripple reactivation (Karaba et al., 2024; McKenzie et al., 2021; Ognjanovski et al., 2017), future studies should test how changes in fine-time scale synchrony depend on these inhibitory plasticity pathways.

### Limitations

The central limitation of our study was that ASV infusion did not indue the expected changes in the ISR. Notwithstanding, the decline in c-Fos resembles that observed following ISR induction where suppression of general translation would be expected to reduce short-lived activity-dependent proteins (Shrestha, Ayata, et al., 2020). Our time point collections may have missed the reported ATF4 and GADD34 peaks at 1 and 3-hrs, which would have been diluted given our small viral infusion volumes, cell-type specific promoter, and bulk hippocampal lysate. Without seeing a coordinated increase in p-eIF2α, ATF4, or GADD34, we cannot attribute the protein changes to the ISR.

Other drugs like anisomycin or cycloheximide have been reported to inhibit protein synthesis by 90-95% (Davis & Squire, 1984). In the previous characterization of the iPKR system, bulk translation was reduced by ∼50% (Shrestha, Ayata, et al., 2020). Future use of an assay like surface sensing of translation (SUnSET) for *in vivo* measurement could provide insight to the degree of translation with our hippocampal manipulation. Additionally, an ASV dose-response curve could isolate whether reactivation of CA1 assemblies was spared because translation was not sufficiently impaired.

Another limitation of our study is that the behavioral task did not include repeated traversals of the same route. Such traversals provide repeated theta sequences (Foster & Wilson, 2006; Diba & Buzsáki, 2007) in which the firing order of CA1 neurons reflects the spatial order of their place fields along the route. A recent study dissociated sequential replay, which depended on entorhinal inputs, from reactivation, which did not (Liu et al., 2023). Therefore, it is possible that changes in CA1 sequencing during replay *do* require CA1 pyramidal plasticity pathway, while patterns of pairwise coincidence do not. Our use of 2D arenas, and relatively low cell yield precluded such an analysis.

## Materials and Methods

### Animals

We used male and female transgenic iPKR (C57/B6/J-Tg iPKR.EGFP fl (Eef2a1)) mice; genotype was verified by tail biopsies sent to Transnetyx (Cordova, TN). Mice were kept on a normal 12-hour light/dark cycle beginning from 6 A.M. to 6 P.M. with water with stable temperature and humidity. Mice were tested during their light cycle. Mice were given food and water *ad libitum*. All animal procedures and breeding were approved by the Institutional Animal Care and Use Committee at University of New Mexico, Albuquerque, NM, USA.

### Viral Cre infusion

Mice were infused with pENN.AAV.CamKII 0.4.Cre.SV40 (AAV5-CaMKIIα-Cre, Addgene #105558-AAV5) or pAAV-CaMKIIα-EGFP (AAV5-CaMKIIα-eGFP, Addgene 50469-AAV5). Glass pipettes were pulled using the Sutter Instrument Co. Model P-97 Micropipette Puller. To deliver the virus, we used the NANOLITER2020 Injector Head (World Precision Instruments) in combination with the MICRO2T SMARTouchTM controller (World Precision Instruments). We delivered 300 nl to dorsal CA1 either unilaterally or bilaterally at a rate of 30 nl/min with a glass pipette back-filled with mineral oil. The coordinates we used from Bregma were AP –2.03, ML +/− 1.5 (or just +1.5 for left unilateral) and from the brain surface DV –1.4 and –1.2 (150 nL per depth). We waited at least 1 minute at the targeted depths before infusing and at least 1 minute before withdrawing.

### Electrode and cannula implantation

We used N = 12 (5 female, 7 male, average age = 217.6 days). iPKR+ mice for electrophysiological recordings. Mice were anesthetized using vaporized isoflurane (1.5% in pure oxygen) implanted with high density probes targeting dorsal CA1 and a cannula targeting the lateral ventricle (from Bregma, AP: – 0.23, ML + 1.0, DV from brain surface –1.5). We used Cambridge Neurotech ASSY 156 P1 (N = 3), 156 F Sharp (N = 2), and 325 D (N = 3) multi-shank electrodes. Cambridge probes were mounted on a movable driver and initially placed 300-400 um from the brain surface before lowering to the left dorsal CA1 (from Bregma, AP –2.03, ML +1.5 from brain surface, 8° angle from the midsagittal plane). A subset of mice (N = 5) were implanted with M1 Precision M-11 linear shank probes directed directly into dorsal CA1 (from Bregma, AP –3.03, ML +1.5, DV –2.00 from brain surface, 25° angle from the midsagittal plane). Upon recumbency from anesthesia, mice were given a 0.1 mg/kg weight of buprenorphine. Mice were allowed post-operative recovery for approximately 7 days prior to beginning baseline recordings.

For mice prepared for Western Blot, we dissected bilateral hippocampi from N = 3 females and N = 3 males iPKR+ mice with bilaterally infused CaMKIIα-Cre and administered ASV for each time point collection (average 124.3 ± 7.3 days old, total N = 6 per timepoint, total N = 30). Control mice were null iPKR mice with the CaMKIIα-Cre virus and received ASV administration collected at timepoint 0 (N = 4). Mice underwent bilateral cannula implantation targeting the lateral ventricles with bilateral guide cannula from Plastics One (26-gauge 2.2 mm center to center, 6 mm depth or 26-gauge 2.0 mm center to center, 2 mm depth, using internal cannula that projected 1 mm past the pedestal).

### Drugs

Asunaprevir (MedChemExpress) was dissolved in DMSO for a stock concentration of 10 mM and diluted in sterile saline to 100 nM. We delivered 2000 nL at 500 nL/min of 100 nM to the lateral ventricle using an injection internal cannula inserted into the stainless-steel guide cannula (Plastics One). Saline and ASV infusions were carried out using an injection cannula extending out of PE50 tubing attached to a 5 uL Hamilton syringe using the UMP3 Microinjection Syringe Pump (World Precision Instruments) with the Micro2T SmartTouchTM controller. The Hamilton syringe was filled with deionized water and the PE50 tubing was filled with mineral oil. The meniscus of saline or ASV was marked prior to infusion to ensure delivery. Upon removal of the dummy cannula either before or after infusion, we checked to ensure there was cerebral spinal fluid coming up from the cannula. Cannula implantation was verified with histology.

### Immunohistochemistry

Mice were transcardially perfused with 1X phosphate-buffered saline (PBS) followed by 4% paraformaldehyde (PFA). Whole brains were dissected and post-fixed in 4% PFA for 3 h, then transferred to 30% sucrose solution until the brains visibly sank. Brains were flash-frozen on crushed dry ice, embedded in optimal cutting temperature (OCT) compound, and coronally cryosectioned at a thickness of 40 µm. Free-floating sections were blocked in 5% goat serum containing 0.1% Triton X-100, followed by incubation in rabbit anti-GFP primary antibody (Thermofisher G10362 1:300) for approximately 72-hrs in 4°C with gentle agitation. Sections were washed with 1X PBS and incubated with either goat anti-rabbit Alexa Fluor 488 or goat anti-rabbit Alexa Fluor 647 secondary antibody (1:200). Following additional PBS washes, sections were counterstained with DAPI for 5-mins and mounted onto Superfrost Plus microscope slides using ProLong Gold Antifade Mountant. Fluorescence labeling was initially assessed using epifluorescence microscopy. Images were subsequently acquired using an Axioscan slide scanner at 20X objective and a Leica TCS SP8 confocal microscope at 63X objective.

### Western Blotting

Hippocampal samples were homogenized and sonicated in 10% weight/volume of homogenization reagent containing Neuronal Protein Extraction Reagent (ThermoFisher 87792) that was supplemented with 10 uL/mg of protease inhibitor (Millipore Sigma P8340) and 10 uL/mg of phosphatase inhibitor (Millipore Sigma P0044) on ice. Protein samples were prepared at 10% (w/v) and the protein concentration was verified using recombinant protein as a reference standard. Samples were run in a pre-cast 4-12% Bis-Tris gels (Invitrogen) and subjected to SDS-Page followed by transfer to PVDF membrane (Bio-Rad) using the Trans-Blot® Turbo™ Transfer System (Bio-Rad). Samples were run with the molecular weight marker PageRuler™ Plus Prestained Protein Ladder (Thermo Scientific 26619). Membranes were blocked in 5% non-fat dry milk or 5% bovine serum albumin in 1X TBS with 0.1% Tween-20 (TBST) and probed overnight at 4° C. Primary antibodies using milk were rabbit anti-PKR (Cell Signaling Technology 3072 1:1000), rabbit anti-GADD34 (Proteintech 10449-1-AP 1:1000), rabbit anti-c-Fos (Cell Signaling Technology 2250 1:1000), and mouse anti-beta-actin (Millipore Sigma A1978 1:10,000). The primary antibody using bovine serum was rabbit anti-ATF (Cell Signaling Technology 11815 1:1000). After washing five times in 1X TBS, membranes were probed with horseradish peroxidase-conjugated secondary IgG goat anti-rabbit (JacksonImmunoResearch Laboratories, Inc 111-035-003) or goat anti-mouse (JacksonImmunoResearch Laboratories, Inc 115-035-003) for 1-hr at room temperature. PVDF membranes were developed using ECL chemiluminescence (Thermo Pierce) using the ChemiDoc MP. Band volumes and Coomassie-stained total lane volumes were quantified by densitometry (Image Lab v 6.1.0 build 7 Standard Edition, Bio-Rad Companies). For total protein normalization, each lane’s Coomassie signal was divided by the mean Coomassie signal of all lanes in the same blot to give a lane normalization factor, and target band volumes were divided by this factor. Values were expressed as fold change relative to the mean of the timepoint 0-hr samples pooled across both sexes. Beta-actin-normalized values (same blot and lane) are shown for comparison. Coomassie lanes outside 0.5-2.0x of the blot reference were flagged but retained. Each lane is an independent sample (n per group = 3 Females, 3 Males, 6 Total).

### In vivo electrophysiology surgical procedures, recordings, and experimental design

Twelve iPKR mice contributed sessions following saline or ASV infusion. An independent reference dataset comprised twelve sessions from three never-infused mice (Huszár et al., 2022); nine of these have annotated context-exposure times and entered the reactivation analyses; the firing-rate and ripple analyses, which need no context marker, used all twelve (84-hrs).

Putative pyramidal cells and interneurons were classified using CellExplorer (Petersen et al., 2021). Sessions comprised 60 min in the home cage, 30 min in familiar familiar context (Familiar 1), a home-cage interval (median 34 min; range 18–134), a 4-min infusion, 30 min in novel context (Novel 1) beginning a median 34 min after infusion onset, 4 h home cage, and subsequent re-exposure to both contexts. Exposures lasted 30 min (range 25–40). Novel 1 began 38 ± 11.8 min after infusion onset. Novel contexts differed across sessions. Every saline and every ASV session with both exposures was analyzed, including repeat infusions; each session’s position in its animal’s infusion sequence was recorded and used as a covariate. ASV sessions nonetheless fall later in the sequence on average, so session order remains partly confounded with treatment. One ASV session (with 10 cells) was excluded from the 2–4 h cell-specific in-ripple estimate because its slope (−1.14) lay 7.4 robust SD from the group median; with it included, the ASV estimate was 0.074 ± 0.084 (p = 0.52).

### Firing and ripple recruitment

Rates were quantified before infusion and 2–8 h afterward, separately by cell class. For the cohort comparison, every iPKR session contributed the period preceding its own infusion, or the whole recording when no infusion occurred (78 session-windows from 12 mice for firing rates; 73 for ripple measures). For neuron *j*, ripple recruitment was the fraction of ripples containing at least one spike. Excess recruitment subtracted the expectation for an independent Poisson process at the neuron’s out-of-ripple rate 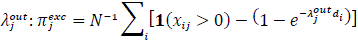, where *d_i_* is ripple duration. Relative gain was the ratio of in-ripple to out-of-ripple firing within the same time window. Rate-based measures were tested on a log_10_ scale. Rates were also computed within wake, NREM and REM using each session’s scored state intervals: spikes in the window and state divided by the time in both, requiring at least 5 min of a state within a window.

Post-infusion rates were scored against each infused session’s own pre-infusion period: pre was the recording from its start to infusion onset (median 2.1 h; range 0.9 to 4.3-hrs) and post was 2 to 8-hrs after infusion (median 5.1-hrs available; each at least 30 min), giving 39 infused sessions from 12 animals, of which two with a single pyramidal unit were not scored. As a robustness check on the infusion comparison, sessions were flagged when the pre-infusion pyramidal median rate was below 0.5 Hz, fewer than five pyramidal units fired both before and 2–8 h after the infusion, or the Spearman correlation of unit rates between these windows was below 0.3 (8 of 39 sessions). The all-state comparison of post-to pre-infusion rates was repeated without them; no other analysis excluded these sessions.

Arms were compared with a permutation test over sessions, a linear mixed-effects model with a random animal intercept (Satterthwaite degrees of freedom), and a sign-flip test over animals with both conditions; the drug effect was also estimated with infusion position (ordinal, or first versus repeat) as a covariate.

### Decomposition of post-experience single-cell changes

For each cell *j*, the change in firing or excess ripple participation between reference and post-experience blocks was modeled as

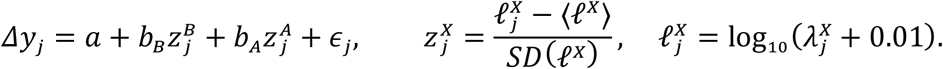

Here 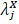 is the firing rate of neuron *j* during running/theta periods of exposure *X*, with *X* = *A* the familiar-context exposure (Familiar 1, drug free) and *X* = *B* the novel-context exposure (Novel 1); 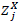 is its within-session *z*-score across pyramidal neurons, so *b_B_* and *b_A_* are changes per standard deviation of activity in each exposure. *a* represents the shared shift in all neurons. *b_B_* represents the change associated with a standard-deviation increase in novel-context activity conditional on familiar-context activity, and *b_A_* the change associated with familiar-context activity conditional on novel-context activity; *b_A_* is a nuisance term that absorbs cell-to-cell differences that were already present before the novel exposure and is not interpreted. The reference block was the home-cage period preceding Novel 1 (drug on board in infused sessions) and the post-experience block the first 30 min after Novel 1; the fit was repeated with the period preceding Familiar 1 (drug free) as reference. The fitted outcomes were log in-ripple rate (offset 0.1), log out-of-ripple rate (offset 0.01), and excess recruitment. Units required at least 10 spikes in each compared block and sessions at least eight eligible units.

A placebo fit used the same model, the same neurons and the same *z*-scores, but took the period preceding Familiar 1 as reference and the period preceding Novel 1 as “post”. Because both blocks end before Novel 1 begins, activity in Novel 1 cannot have caused any change between them, so a nonzero *b_B_* there reflects slow drift in excitability (cells rising over the session are low in Familiar 1 and high in Novel 1, predicting *b_B_* > 0 and *b_A_* < 0). The real-minus-placebo difference was computed within session, using the fit with the period preceding Familiar 1 as reference, and tested by sign-flip over sessions.

### Pairwise reactivation

Exploration templates comprised cell-pair spike-count correlations during running or theta in 100-ms bins. Ripple co-activity was measured in home-cage periods before (PRE) and after (POST) each exposure. All pairs of scored units were included. With *r_pre_* and *r_post_* denoting the correlations of the template’s pairwise-correlation vector with PRE and POST vectors, and *r_pp_* the correlation between PRE and POST vectors, directional reactivation was calculated as

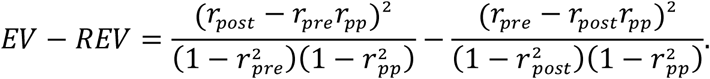

POST was up to 30 min after exposure except in explicitly indicated later windows; for Familiar 1 it ended at infusion onset and was shorter than 30 min in 12 of 35 exposures (18–30 min; seven ASV, five saline). For Novel 1, PRE was the intervening pre-exposure home-cage period excluding the 30 min after infusion onset (22–134 min, median 39); for other exposures it was the preceding home-cage hour. The 30-min exclusion was timed from infusion onset, about 26 min after the 4-min infusion ended, so Novel 1 PRE comprised pre-infusion home cage plus 0–10 min of post-infusion home cage. Each template required at least 10 units with at least 100 spikes each. Rate asymmetry between blocks was assessed by binomial thinning (20 draws).

### Pair coincidence model

For ripple *i* of duration *d_i_* and pyramidal neuron *j*, spike counts *x_ij_* were fitted by a Poisson model,

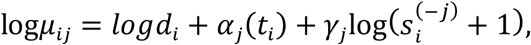

where *α_j_* takes one level per 30-min block (ridge-penalized), 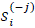 is the summed count of the other recorded pyramidal neurons, and *γ_j_* is estimated separately for ripples before and after the exposure. Coincidences *y_ijk_* = *x_ij_x_ik_* were modeled with offset log(*μ_ij_μ_ik_*), recomputed with *s_i_* excluding both neurons, as

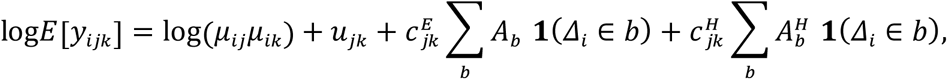

where *Δ_i_* is the time after the end of the exposure in hours, *b* indexes 1-h bins after the exposure (0–1, 1– 2, 2–3 and 3–4 h), 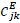 the partial correlation of the pair’s 100-ms spike counts during running or theta epochs of the exposure with the population count partialled out, 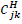 the same quantity from a duration-matched home-cage block outside the fitted range, both *z*-scored across pairs, and *u_jk_* a pair fixed effect removed by conditioning on each pair’s total coincidences. Pre-exposure ripples carry no bin term, so their level is absorbed by *u_jk_*: *A_b_* is the excess in bin *b* relative to the pre-exposure ripples, per standard deviation of *c^E^*, and 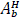 the same for the home-cage template, so that *A_b_* is the exploration-specific excess beyond any change in home-cage pair structure. No decay form was assumed. Home-cage ripples within 4-hrs of the exposure were used, excluding the 30 min after infusion onset; for an exposure before the infusion, the post-exposure window ended at the infusion. Neurons required at least 20 spikes during exploration and 10 ripple spikes on each side, and pairs were capped at 1,500 per session. A session contributed to a bin only if at least 20 post-exposure ripples fell in it.

To locate the latest time at which the excess remained detectable, the model was refitted with 15-min bins, with the home-cage template entered as a single post-exposure term and a drift term *δ*|*Δ_i_*| symmetric about the exposure added to the exploration template (*δ* = 0 in the primary fit; estimated in a sensitivity analysis). Bins were pooled within animal by their model information. Starting from the first bin, a window was extended one bin at a time until its animal-level excess was positive at sign-flip p < 0.05, then closed, and the next window began after it. Because extending a window until it passes inflates its nominal p, each window starting at bin a was recalibrated: the statistic was the maximum t over all windows beginning at a, and its null distribution was obtained by flipping each unit’s sign jointly for all bins from a onward. In simulation this procedure held the false-positive rate at 0.045 with an early-only effect detected in 98% of runs. The latest window with calibrated p < 0.05 is reported.

### Coordination across exposures

For each of the four exposures (Familiar 1, Novel 1, Familiar 2, Novel 2), spike counts of pyramidal neurons were binned at 100-ms within running epochs and divided into 1-s windows. For each neuron, the summed count of the other recorded pyramidal neurons was regressed out, and pairwise coordination 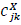 was the correlation of the residuals of neurons *j* and *k* in exposure *X*. Neurons required at least 20 spikes in each of the four exposures. Similarity between exposures *X* and *Y* was *z*(*X*, *Y*) = 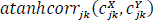. The template contrast was *z*(*Novel* 1, *Novel* 2) − *z*(*Familiar* 1, *Novel* 2). Because Novel 1 is closer in time to Novel 2 than Familiar 1 is, the primary estimate came from a per-session fit over the six exposure pairs,

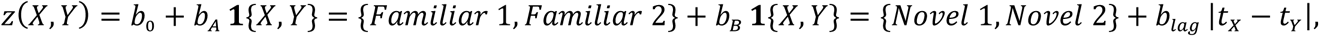

where *t* is the midpoint of an exposure in hours and *b_B_* is the re-expression of novel-environment coordination beyond elapsed time. Split-half reliability of *c^X^* was obtained from alternating 1-s windows (Spearman–Brown corrected), and similarities were also computed after dividing each correlation by the square root of the two reliabilities. Sessions required all four exposures in the order Familiar 1, Novel 1, Familiar 2, Novel 2 and at least five neurons meeting the spike criterion. Arms were compared as described below, and again with days since the animal’s first recording day, or the number of earlier recording days, as a covariate.

### Statistical analysis

The animal was the unit of replication for paired treatment contrasts and for within-arm tests, which used animal means. The session was the unit for across-session associations where linear mixed-effects models included treatment as a fixed effect with animal and session random effects and Satterthwaite degrees of freedom. To compare regressors against zero, exact two-sided sign-flip tests were applied to paired and within-arm effects, with a floor of 2/2*^m^* for *m* animals. Drug effects on time-varying outcomes were further assessed as within-animal differences in the pre-to-post change between ASV and saline days. Across-session associations used Spearman rank correlation. Sample sizes varied with eligibility criteria. Comparisons involving never-infused recordings used the session as the unit of replication (nine sessions from three mice with annotated context times; twelve sessions for rate and ripple measures).

### Use of Generative AI

ChatGPT and Claude were used during manuscript preparation to design analyses and figures and structure the initial outlines. The authors subsequently revised, rewrote, and critically edited the AI-assisted material, ensuring scientific accuracy and appropriate interpretation.

## Data availability

In vivo electrophysiology data (DANDI ID 002096)

Full Western blots available on Mendeley (Yang, Gaokhia (2026), “Disrupting translation in CA1 pyramidal cells blocks novelty-induced increases in population firing rate while preserving newly formed assemblie”, Mendeley Data, V1, doi: 10.17632/zb5g77c4×3.1)

## Software availability

All software can be found at https://github.com/McKenzieNeuro/McKenzieLab/tree/main/transPlasticity/final_figs. runAllAnalyses.m performs every analysis of the electrophysiological data and makePaperFigures.m plots every figure.

## Author contributions

G.Y. collected electrophysiology data, processed histology, conducted Western blotting and contributed to writing the manuscript. N.M. generated the tissue needed for Western Blotting and contributed to developing protocols for protein analysis. B.T.J. helped with electrophysiology data collection and histology. M.B. contributed to histology and Western blotting. N.P.B assisted in experimental design and protein analysis. S.M. performed initial experiments, analyzed data, and contributed to writing the manuscript.

## Acknowledgements

We thank Dr. Joseph Kunkel for assistance in database management. iPKR mice were a gift from Dr. Eric Klann and Dr. Prerana Shrestha. S.M. was supported by NIMH R01MH140076, NIMH R00 MH118423, NINDS RC2NS138557, and a Whitehall Foundation Research Grant. G.Y was supported by NIH/NIGMS T32 GM144834. This work was selected as a winner for the Hidden Science Competition hosted by Astera Institute (https://ror.org/00ydx1s47).

---

**Supplemental Figure 1.**
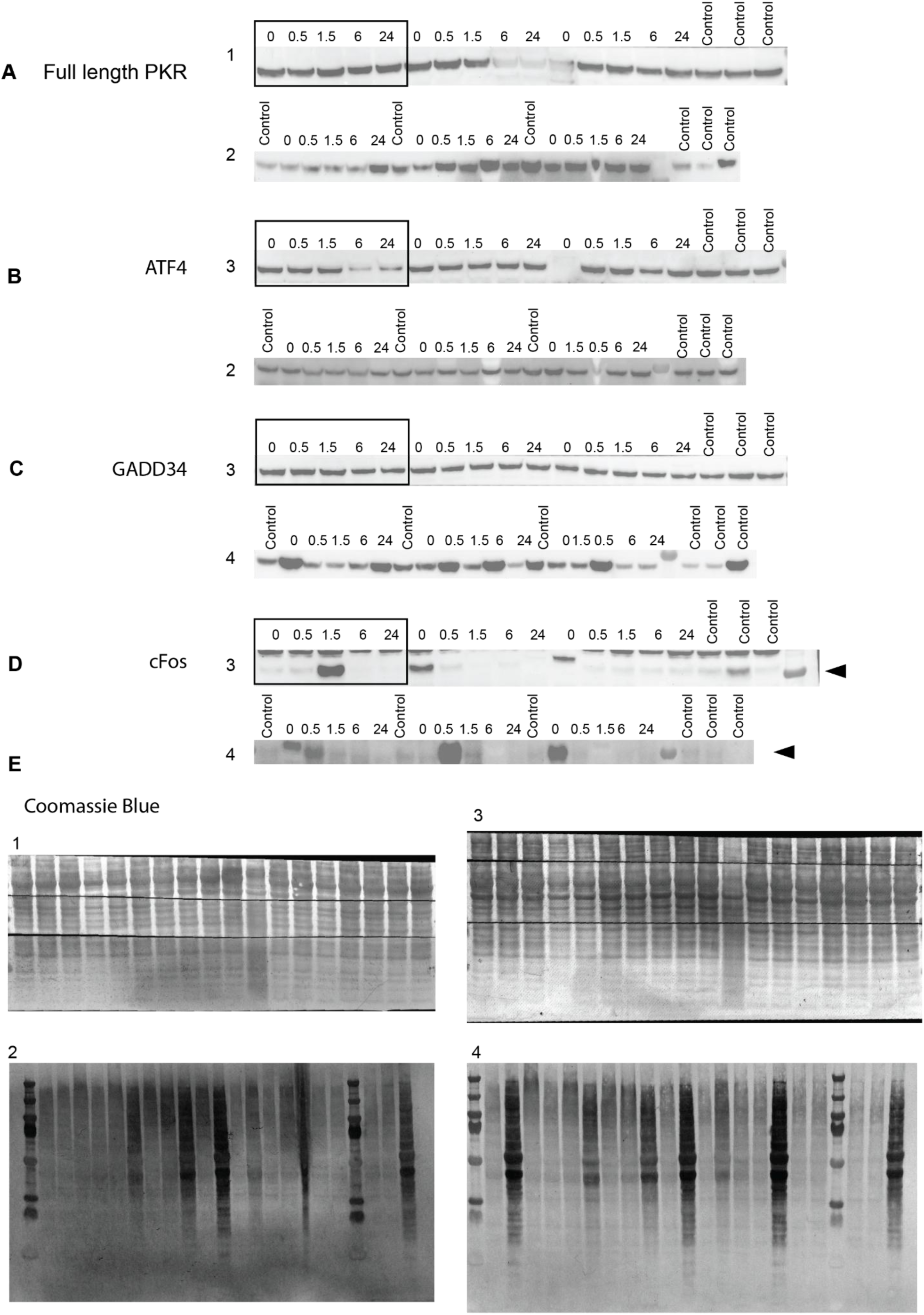
Full Western blot panel. Cut blots shown for **A,** full length iPKR using the PKR antibody from Cell Signaling Technology, **B,** ATF4. **C,** GADD34, **D,** cFos, and **E,** Coomassie staining after probing for antibodies per blot for total protein normalization. Black squares indicate the examples used for the primary example in figure 2. Black arrow indicates which band was quantified.

**Supplemental Figure 2.**
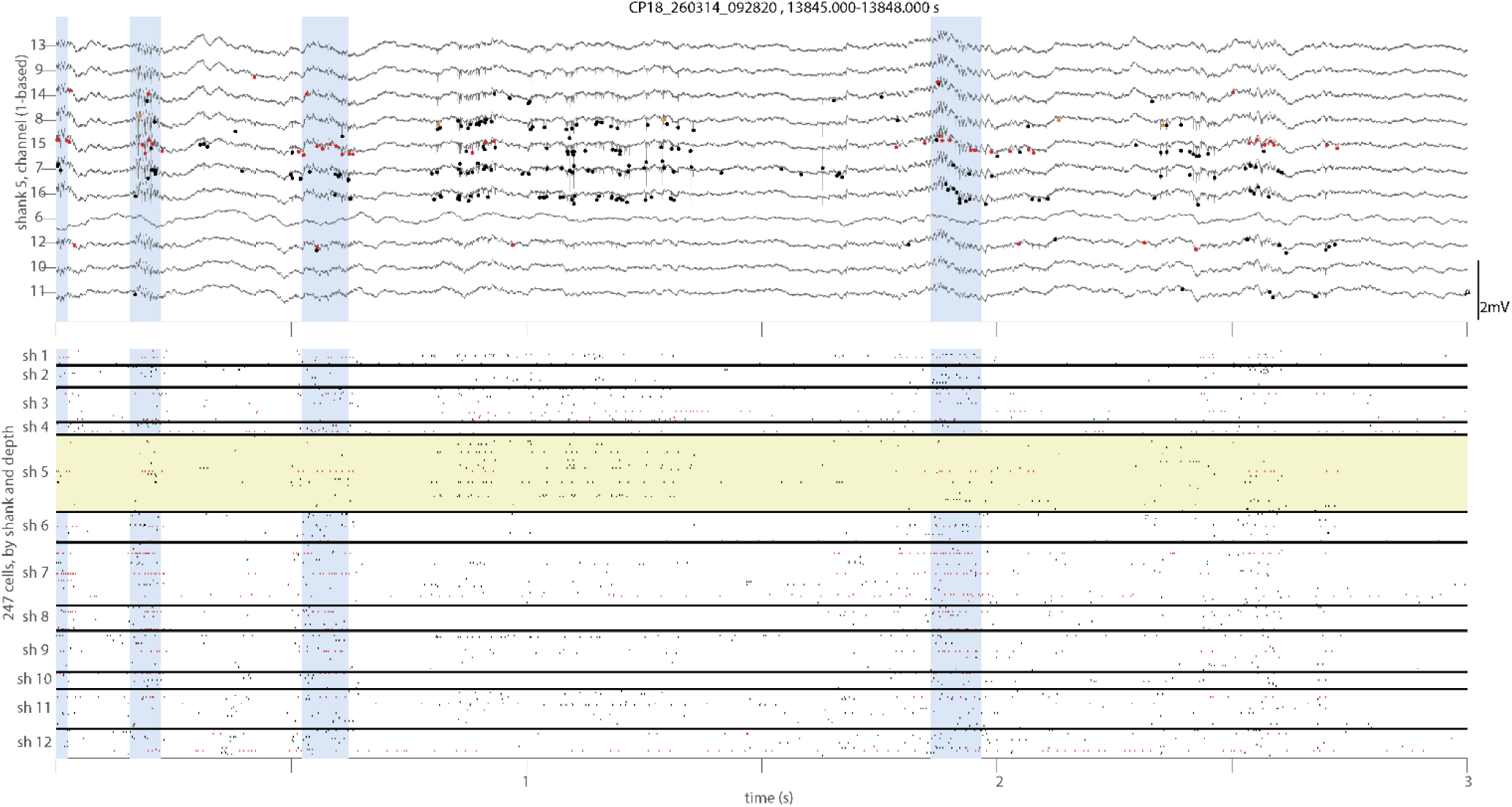
Example recording of wideband signals and population spiking. Three seconds of data from one session (CP18_260314_092820, 13,845–13,848 s). Top, wideband signals (30 kHz) from the 11 channels of one shank (shank 5), ordered by position along the shank. Dots mark the spikes of cells on this shank on each cell’s peak channel, colored by putative cell type (black, pyramidal cells; red, narrow-waveform interneurons; orange, wide-waveform interneurons). Scale bar, 2mV. Bottom, spike rasters of all 247 cells recorded across 12 shanks, grouped by shank and ordered by depth within each shank, with the same colors. Note high baseline activity between ripples. Rows of shank 5 are shaded yellow. Blue shading marks detected sharp-wave ripples.

**Supplemental Figure 3.**
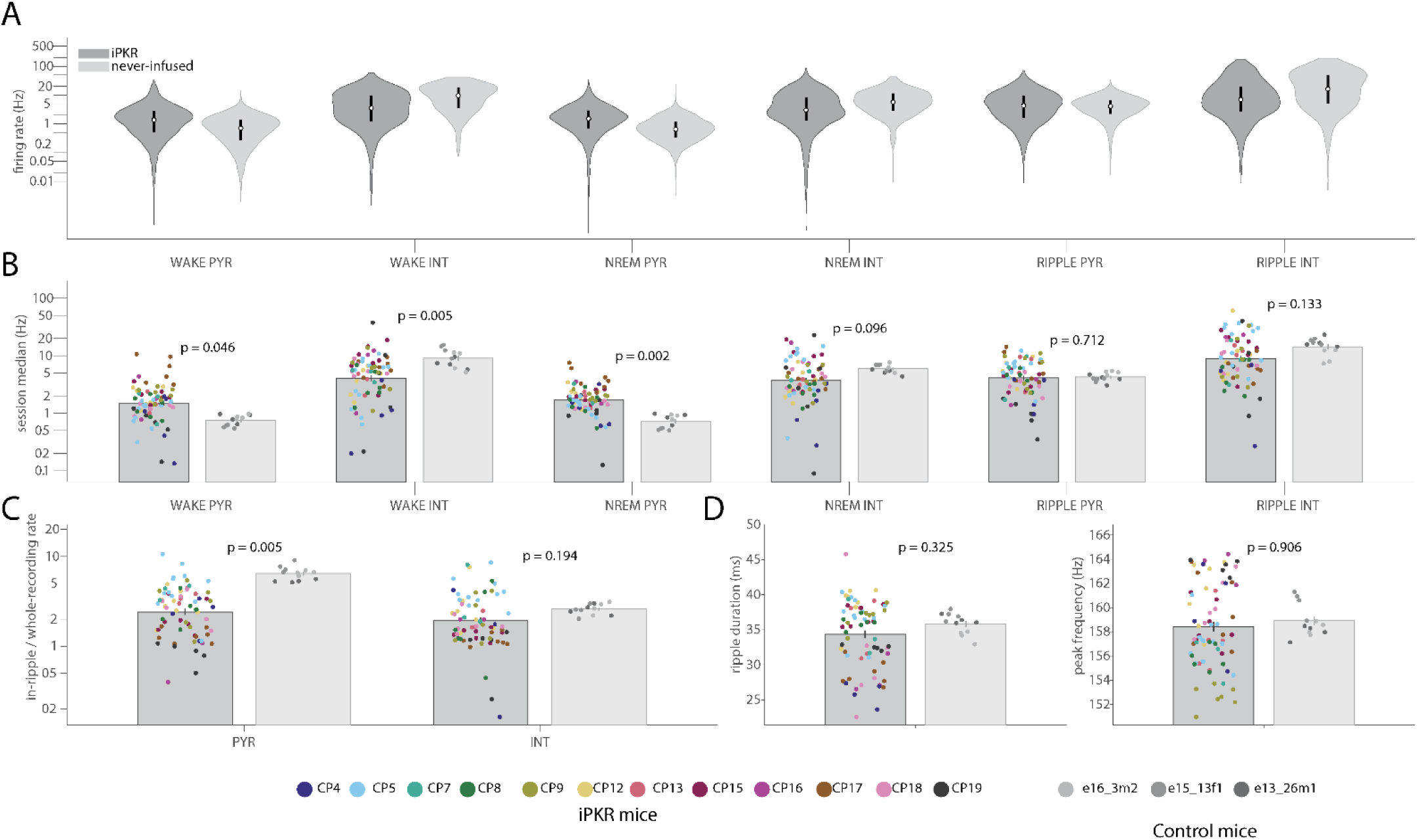
Baseline CA1 firing and ripple properties differ between iPKR and never-infused mice. ***A***, Distributions of firing rates of individual pyramidal cells (PYR) and interneurons (INT) during wake, NREM sleep, and sharp-wave ripples in iPKR (dark) and never-infused (light) mice, plotted on a log scale. Each distribution pools cells across sessions; cells silent in a given state are omitted. Cell counts (iPKR/never-infused) were: wake PYR, 2904/2544; wake INT, 1392/491; NREM PYR, 2909/2382; NREM INT, 1429/453; ripple PYR, 2793/2543; ripple INT, 1246/490. ***B***, The same measures summarized per session; each dot is the median rate of one session, colored by mouse. Session counts (iPKR/never-infused) were: wake PYR, 72/12; wake INT, 70/12; NREM PYR, 72/11; NREM INT, 71/11; ripple PYR, 71/12; ripple INT, 68/12. ***C***, Ripple gain (in-ripple firing rate divided by whole-recording rate) for PYR (71/12 sessions) and INT (68/12) on a log scale. **D,** ripple duration (geometric mean; 73/12), and ripple peak frequency (73/12). Bars in ***B*** and ***C*** show mean ± SEM across sessions. iPKR measures were taken before each session’s infusion, or across the whole recording in sessions without one. *p* values compare groups using a linear mixed-effects model on session values with a random intercept for animal (Satterthwaite degrees of freedom).

